# Modeling three-dimensional structures of flaviviral exoribonuclease-resistant RNAs by combining functional and computational analyses

**DOI:** 10.64898/2026.11.24.769036

**Authors:** Daniiar Zhaguparov, Sneha Munshi, Rohith Vedhthaanth Sekar, Ishrat Meghla, Michael T. Wood-side

## Abstract

Exoribonuclease-resistant RNAs (xrRNAs) are non-coding RNAs used by viruses to evade host RNase-mediated degradation. In flaviviruses, xrRNAs are thought to share an unusual knot-like fold with the 5′-end threaded through a ring formed by a multi-helix junction and closed by a pseudoknot, but only a handful of high-resolution structures have been solved to date. We combined structure prediction tools with mutational analysis of RNase resistance to determine previously unsolved 3D structures of xrRNAs from the West Nile virus (WNV) and two dengue virus (DENV) serotypes. We first predicted possible pseudoknot structures computationally, then tested them by systematically mutating putatively pseudoknotted bases and assessing RNase resistance to discriminate between models. The successful pseudoknot models were used to predict 3D structures computationally, refining the models by identifying tertiary contacts at the threaded 5′ end via another round of mutagenesis and RNase resistance testing. We found that all 3 xrRNAs shared the same threaded knot-like architecture as in experimentally solved flaviviral xrRNAs, with similar tertiary contacts generating an analogous hierarchy of interactions importance for RNase resistance, but the pseudoknot length varied considerably, from 8 pairs in WNV to 2 in DENV2. This work illustrates a powerful new approach for determining xrRNA structures that are hard to obtain by high-resolution experimental methods by integrating computational predictions with functional assays.

## INTRODUCTION

Exoribonuclease-resistant RNAs (xrRNAs) are structured RNA elements found in the genome of many positive-sense single-stranded RNA viruses [1-7]. These elements play a crucial role in viral metabolism by stalling the progression of a wide range of 5′→3′ directional host exoribonucleases, thereby protecting the downstream viral sequences from degradation [1, 2, 8-10]. In flaviviruses (a family of positive-sense RNA viruses including Zika, West Nile, dengue and many others that pose a threat to human populations worldwide [11, 12]), xrRNAs are often located in the 3′ untranslated region (UTR) [1, 3, 8]. As a result, incomplete 5′-to-3′ degradation of the viral genome mediated by cytoplasmic RNases, such as the highly processive enzyme Xrn1 [13-15], leads to the production of non-coding RNA fragments of different lengths [1, 2, 8]. These subgenomic fragments are functional, acting to enhance viral cyto-pathicity and pathogenicity [16-19] by dysregulating antiviral responses through various processes such as inhibiting interferon signalling [2, 20, 21], impairing mRNA turnover [8], and supressing siRNA and miRNA production in host cells [22, 23]. Beyond flaviviruses, xrRNA elements have been identified in a wide range of evolutionary distant viral families [24-27], including plant-infecting viruses [4-7, 9] and even negative-sense phleboviruses and ambisense arenoviruses [28], suggesting that they form a class of RNAs that are generally important for viral lifecycles across major viral families. xrRNAs thus offer a potential new target for antiviral therapeutics for currently untreatable viral infections.

Phylogenetic analysis has found xrRNA sequences in all genera of *Flaviviridae* [27], suggesting xrRNAs play an important role in such viruses. Covariance modeling of flaviviral xrRNA sequences was used to suggest the presence of two distinct classes based on their proposed secondary structures [10], but more recent studies have argued all flaviviral xrRNAs share the same basic fold architecture [24, 25, 27, 31]. Only a handful of high-resolution 3D structures of flavivirus xrRNAs have been solved to date, however [1, 29-31]. Intriguingly, all exhibit the same, unusual knot-like topology, wherein the 5′-end is threaded through a ring formed by a multi-helix junction and closed by a pseudoknot (PK) interaction (Fig. 1A), a topology otherwise seen to date only in coronavirus frameshift-stimulatory pseudoknots [32-37]. Analysis of flaviviral xrRNA sequences as well as low-resolution structural probes suggest that these xrRNAs differ from one another mainly in the pseudoknot length, ring size, presence/absence of unpaired conserved nucleotides in the helical junction, and tertiary contacts anchoring the 5′-end (Fig. 1B) [3, 24, 25, 27, 38]. However, the 3D structures of most such xrRNAs remain unknown.

**Fig. 1:**
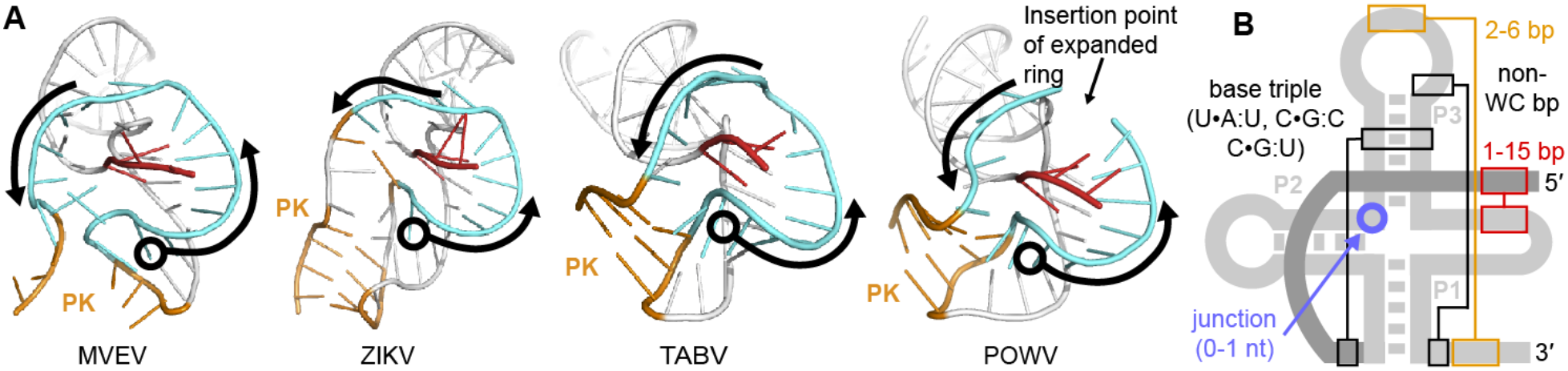
Knot-like topology of flaviviral xrRNAs. Topology with the intramolecular ring (cyan) being formed around the 5′-end (red) and closed by a pseudoknot (PK, orange) is observed in crystal structures of (A) Murray Valley Encephalitis virus xrRNA2 (PDB ID 4PQV) [1], Zika virus xrRNA1 (PDB ID 5TPY) [29], Tamana bat virus xrRNA (PDB ID 7K16) [30], and Powassan virus xrRNA1 (PDB ID 9DP9) [31] ring-knots. (B) Cartoon representation of a general secondary structure of the functional core in *Flaviviridae* xrRNAs with highlighted regions proven to be important for RNase resistance.

Although the knot-like topology in the functional core of flaviviral xrRNAs was originally proposed to be responsible for their RNase resistance, because the ring closed by the PK wraps around the 5′-end presumably blocks translocation of the enzyme [29], studies of the Zika virus (ZIKV) xrRNA showed that the critical element for generating RNase resistance is not the topology of the fold but rather the fact that the xrRNA is extremely resistant to mechanical unfolding by the enzyme [39, 40]. This mechanical resistance arose from key tertiary contacts: the pseudoknot closing the ring around the threaded 5′ end and interactions between the ring and the 5′ end that hold the end rigidly [40]. Deleting these contacts led to a loss of RNase resistance proportional to the inability to form a mechanically resistant structure [39, 40]. These observations suggest that it should be possible to model flaviviral xrRNA structures by using point mutants to identify the contacts generating RNase resistance and then combine these observations with computational modeling to build 3D structures.

Here we have applied this approach to obtain 3D structural models of three flaviviral xrRNAs whose structures remain unsolved: one from West Nile virus (WNV), which is the most common mosquitoborne virus in North America [41], and two sero-types of dengue virus (DENV), DENV2 and DENV4, which is the most prevalent mosquito-borne virus worldwide [42]. Iterating between computation and experiment, we first used sequence-based predictions of potential pseudoknot structures to guide the choice of point mutants for study in RNase-resistance assays. The pseudoknot structures identified by the enzymatic assays were then modeled in 3D computationally, using additional mutagenesis and RNase-resistance assays to identify tertiary contacts on the 5′ end for refining the structural model. The structures obtained this way all showed the same knot-like topology (featuring a ring surrounding the 5′ end held in place by tertiary contacts) as previously observed in high-resolution experimental structures of flaviviral xrRNAs [1, 29-31], supporting the universality of the self-threaded architecture, but they had pseudoknots varying in length from 2 to 8 base-pairs.

## MATERIALS AND METHODS

### Secondary structure predictions

Secondary structures of previously identified xrRNA1 sequences from WNV (RefSeq NC_001563.2) and two lineages of DENV (DENV2: RefSeq NC_001474.2, DENV4: RefSeq NC_002640.1) were predicted by using four different platforms able to predict pseudoknot formation: HotKnots [43, 44], pKiss [45], pKnots [46], and Nupack [47, 48]. Because the first four nucleotides in the 5′ end of the xrRNAs are expected from phylogenetic studies [10, 24, 25] to be involved in complex tertiary contacts, they were excluded from the secondary structure predictions.

### RNA synthesis

DNA inserts encoding WNV, DENV2 and DENV4 xrRNAs were cloned into the pcDNA5/FRT/TO vector (Addgene) between Bam-HI and HindIII sites. DNA transcription templates were prepared from the resulting plasmids using PCR, containing a T7 promoter sequence in the upstream primer, a 56-bp-long leader sequence, xrRNA sequence, and a 29-bp-long downstream sequence for Xrn1 digestion assay *in vitro*. For all xrRNAs, mutations were either introduced into the template for the wild-type xrRNAs by site-directed mutagenesis, or by directly cloning DNA inserts encoding desired mutations. Templates were transcribed *in vitro* by T7 RNA polymerase to generate RNA transcripts used in measurements. Resulting RNA sequences are listed in Table S1. To generate RNA with a monophosphate group at the 5′-end as needed for loading Xrn1 enzyme, 1 μg of RNA transcript was incubated with pyrophosphohydrolase (RppH, New England Biolabs) for 2 h at 37°C following the manufacturer′s protocol. The reaction was terminated by adding 1 µL of 500 mM EDTA and heating the sample to 65°C for 5 min. In preparation for digestion assays, roughly 200 ng of the RNA product was heated at 90°C for 3 min and then cooled at room temperature for 5 min to refold into the native structure. Finally, 1 µL murine RNase inhibitor (40 U/μL, New England Biolabs) was added to prevent nonspecific degradation.

### Xrn1 digestion assay

Roughly 100 ng of refolded xrRNA construct was placed in each of two tubes, and eiether1 µL of Xrn1 (1 U/µL, New England Biolabs) or 1µL of RNase-free water was added to the tube, respectively for (+) and (−) Xrn1 measurements. Reactions were incubated for 2 h at 37°C and terminated by adding 5 μL of formamide loading buffer, heating to 90°C for 3 min, and loading the products into an 8% denaturing polyacrylamide gel for gel electrophoresis. Gels were stained with SYBR Green II (Invitrogen) for 15 min before imaging. The average percentage of exoribonuclease resistance (XR) was quantified using ImageJ v.1.8.0 [49], averaging over three or more gels from biological replicates.

### Molecular dynamics simulations

After confirmation of the secondary structure from the experimental tests, the 3D structures of xrRNA molecules were predicted using Rosetta FARFAR2 [50] and/or Sim-RNAweb2.0 [51], using the latter when specific tertiary contacts like base triples needed to be enforced. Structural modelling and subsequent assessment of conformational stability using all-atom molecular dynamics (MD) simulations followed a previously described workflow [52]. Briefly, for each xrRNA sequence, the proposed secondary structure was used as input to the generate 3D models. The lowestenergy cluster centre models from the 3D structure predictors were inspected, and the lowest-energy threaded conformation was selected for MD simulation studies. All-atom MD simulations of xrRNA molecules were performed using Amber24 [53].

Each RNA was solvated in an OPC explicit-water box with a minimum solute–box margin of 12 Å and parameterized using the OL3 RNA force field. Systems were neutralized with Na^+^ counterions, and additional Na^+^ and Cl^−^ ions were added to reach 0.15 M NaCl. Because Mg^2+^ ions are known to play an important role in xrRNA folding and mechanical stability [39], separate simulations were also run including Mg^2+^ ions, where the ions were manually positioned based on the ZIKV xrRNA crystal structure (PDB ID: 5TPY) by structurally aligning 5TPY with the predicted xrRNA models and transferring the corresponding ion positions. All simulations were carried out at 310 K. For each construct and condition, three independent 1-µs trajectories were generated. To obtain representative conformations, the final 500 ns of each replicate were pooled for clustering analysis using an RMSD cutoff of 3 Å, calculated over pseudoknot atoms excluding highly flexible stem loops P2 and P4; the centroid of the most populated cluster was taken as the representative structure for the xrRNAs.

## RESULTS

To build atomistic models of the structure of WNV, DENV2, and DENV4 xrRNAs, we iterated computational predictions with experimental tests and refinement as illustrated in Fig. 2. We first used four different secondary structure solvers, all able to predict the formation of RNA pseudoknots (Hot-Knots [43, 44], pKiss [45], pKnots [46], and Nupack [47, 48]), to identify potential pseudoknot regions within the selected xrRNA sequences (step 1). We then created a series of mutants containing point mutations chosen to disrupt the predicted PK regions (step 2), and performed Xrn1 digestion assays on the mutants to assess which pseudoknots remained intact despite the mutations, as assessed from any change in the exoribonuclease resistance (XR) of the xrR-NAs (step 3). Discarding all putative secondary structures that were inconsistent with XR results for the mutants (step 4), we used the remaining secondary structures to predict the 3D structure using Rosetta FARFAR2 [50], and took the lowest-energy prediction as the input for 1-μs all-atom molecular dynamics (MD) simulations (step 5), to obtain a single thermodynamically stable 3D structural model for each xrRNA. Since the secondary structure prediction did not account for tertiary contacts expected at the 5′ end [54], we next made additional point mutations in the 5′ end (step 6) and tested their effects on XR levels (step 7), refining the final structural model with the tertiary contacts identified in this way (step 8).

**Fig. 2:**
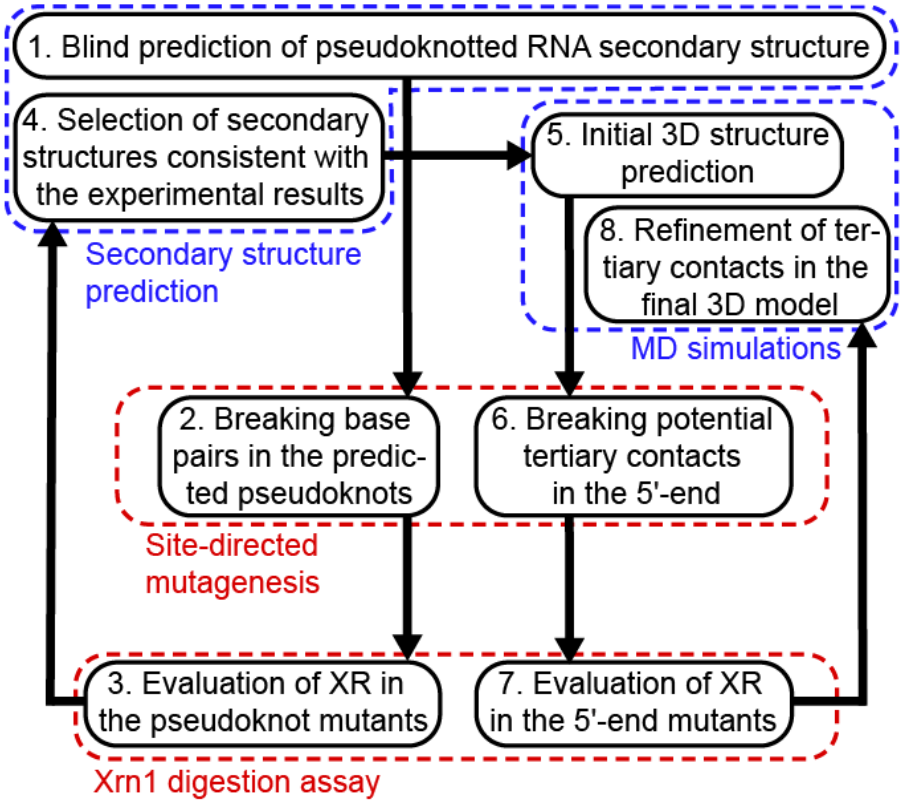
Iterative workflow used to obtain 3D structural models of flaviviral xrRNAs. First, the potential pseu-doknot regions are identified in blind secondary structure predictions obtained by using HotKnots [43, 44], pKiss [45], pKnots [46], and Nupack [47, 48] software using xrRNA sequences. Second, base pairs in the predicted pseudoknots were disrupted by site-directed mutagenesis and RNase resistance of these mutants was tested in Xrn1 digestion assay. Next, secondary structures consistent with the mutational analysis were used as an input for initial 3D structure predictions. Lastly, tertiary contacts in the 5′-end, which were proven experimentally through mutagenesis and enzymatic digestion assay, were added to the final 3D models refined by molecular dynamics. Computational and experimental approaches are shown in blue and red boxes, respectively.

### Modeling the WNV xrRNA1 structure

We first applied this procedure to the first of the xrRNAs in WNV. Predicting pseudoknotted secondary structures from the sequence (Fig. 3A), we found that 3 of the 4 predictors yielded almost identical base-pairing, which generally followed the 2D architecture suggested by previous phylogenetic work [24, 25, 27]. However, one predictor (pKiss) favored a different fold featuring a double pseudoknot. To distinguish between these two folds, we mutated six nucleotides (G35C/A36U/G56C/A60U/A61U/C62G), which should leave the double-pseudoknot intact but break base pairs in the single pseudoknot (Fig. 3A, blue), such that RNase resistance should be unchanged if the double-pseudoknot is present but reduced or eliminated if the single pseudoknot forms. Enzymatic measurements of XR levels (Fig. 3B) found that XR dropped from 86 ± 3% in the wild type (WT) to 57 ± 8% in this mutant, ruling out the double-pseudoknot fold. Notably, the persistence of significant XR even after breaking 6 pairs in the single pseudoknot also contradicts a previous proposal that the pseudoknot consists of only 6 pairs [38, 55, 56], 5 of which would have been broken in this mutant. To distinguish between the 7-bp and 8-bp pseudoknot models, we inserted an additional mutation, G40C, which should disrupt the extra pair in the model with the 8-bp pseudoknot but not further affect the 7-bp pseudoknot (Fig. 3B, green). Given that this mutation completely abolished XR (0 ± 2%), the 8-bp pseudoknot model (Fig. 3A, inset iii) was selected as the basis for modeling the 3D structure in FARFAR2.

**Fig. 3:**
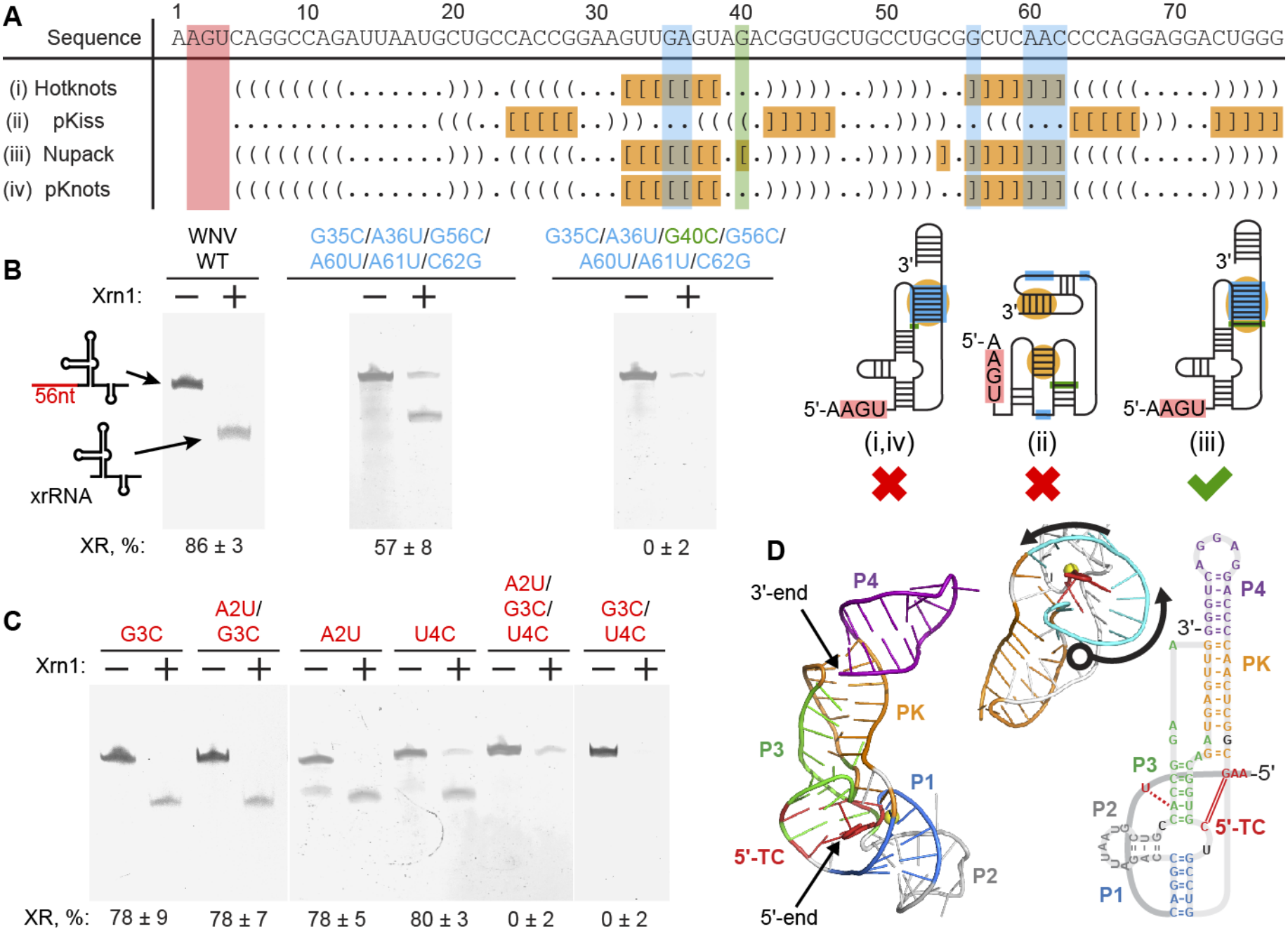
Modeling 3D structure of WNV xrRNA1. (A) Sequence and potential secondary structures predicted by four structure solvers, shown in dot-bracket notation and cartoons (insets i-iv). Orange: predicted pseudoknot areas; blue: nucleotides mutated to target pseudoknot interactions; red: nucleotides mutated to target 5′-end tertiary contacts. (B) Representative gels of Xrn1 digestion for pseudoknot mutants. Adding Xrn1 produces resistant fragment (xrRNA) or fully digested RNA (if no resistance). XR activity calculated from fraction of RNA remaining undigested. Secondary structure consistent with RNase digestion marked with green checkmark (panel A inset). (C) Representative gels showing digestion of 5′-TC mutants. (D) Final 3D structural model and 2D representation with colour-coded stem-loops P1-P4, pseudoknot (PK), 5′-end tertiary interactions (5′-TC), and Mg^2+^ ion (yellow). Final structure features knot-like fold with 5′-end (red) threaded through intramolecular ring (cyan) closed by pseudoknot (orange).

Taking the lowest-energy 3D structure prediction from FARFAR2, we ran 1-μs all-atom MD simulations to obtain a better model of the equilibrated structure. The G40:C54 pair was not persistent in the MD simulation, but since its existence was established by the mutagenesis, we constrained it to stay intact by applying heavy positional restraints on its atoms in MD simulations. The most populated cluster in the equilibrated MD (Fig. S1A) showed the same knot-like architecture seen in the existing crystal structures of flaviviral xrRNAs, with the 5′ end threaded through a ring closed by the pseudoknot.

The 5′ tertiary contact (5′-TC) G3:C47 was formed in the simulations, even though this pair was not included in the input to FARFAR2. However, this model lacked two other stabilising tertiary interactions in the 5′ end that have been suggested by phylogenetic analysis: the triple U4•A25:U45 and the pair A2:U48 [10, 24, 25, 27]. Such 5′-end stabilizing tertiary interactions are proposed to be crucial for xrRNA function [29, 39, 40, 55].

To test if these 5′-end interactions are truly absent or just escaped prediction in the modeling, we generated new mutants disrupting various combinations of the proposed contacts (Fig. 3A, red) and tested them enzymatically for XR (Fig. 3C). Mutating U4, G3, or A2 individually to break the proposed contacts did not change XR levels, nor did mutating A2 and G3 simultaneously, but mutating both G3 and U4 abolished XR, as did mutating all 3 bases simultaneously. These results suggest that A2:U48 is not present or plays no role in RNase resistance, but that U4 does form the proposed tertiary contact. To incorporate the U4•A25:U45 triple into the 3D model, we used SimRNAweb v.2.0 platform [51] to place a hard pairing restraint on U4•A25 while including the A25:U45 pair in the secondary structure restraint.

Simulations were performed using the top cluster centre with threaded topology with and without Mg2+ ion placed at the same position as in ZIKV xrRNA1 crystal structure [29]. Notably, the tertiary contacts persisted only in the simulations containing a Mg^2+^ ion, suggesting this ion plays an important role in the stability of the 5′ end contacts. This observation is consistent with previous work on the ZIKV xrRNA [39]. The final model, consisting of the most-populated cluster in these simulations, is shown in Fig. 3D; an overlay of the top 3 cluster centres with RMSD and RMSF plots is shown in Fig S2A.

### Modeling the DENV4 xrRNA structure

We next studied the xrRNA from DENV4, which is unusual because DENV4 is thought to have a single xrRNA structure instead of the multiple xrRNA elements found in most other flaviviruses, including the other DENV serotypes [57, 58]. Such a difference might suggest the single DENV4 xrRNA differs in some key ways from those of other flaviviruses, making it particularly interesting. Secondary structure predictions (Fig. 4A) suggested two different double-pseudoknot folds and two similar singlepseudoknot folds (Fig. 4A, insets i–iv). To distinguish these structures, we first mutated bases G48 or G48 and G29, to break base-pairs in one of the predicted pseudoknots (Fig. 4A, purple). The single mutation G48C halved XR, from 79 ± 5 % in WT to 45 ± 7 % in the mutant, whereas the double mutation G29C/G48C completely abolished XR (Fig. 4B). In contrast, mutations to G15, G62, and/or G64 designed to destabilize one of the other predicted pseudoknots had no effect on XR levels. Taken together, these results imply the presence of a single 4-bp pseudoknot, consistent with previous phylogenetic analysis [55, 56]. However, our analysis could not distinguish between the two very similar variants of the single-pseudoknot fold, which differed in the presence of an extra 2-bp stem in a region that was previously reported not to contribute to XR [29, 59]. We therefore used both secondary structure models as inputs to 3D predictions by FARFAR2 so that the stability of their interactions in 3D structures could be tested.

**Fig. 4:**
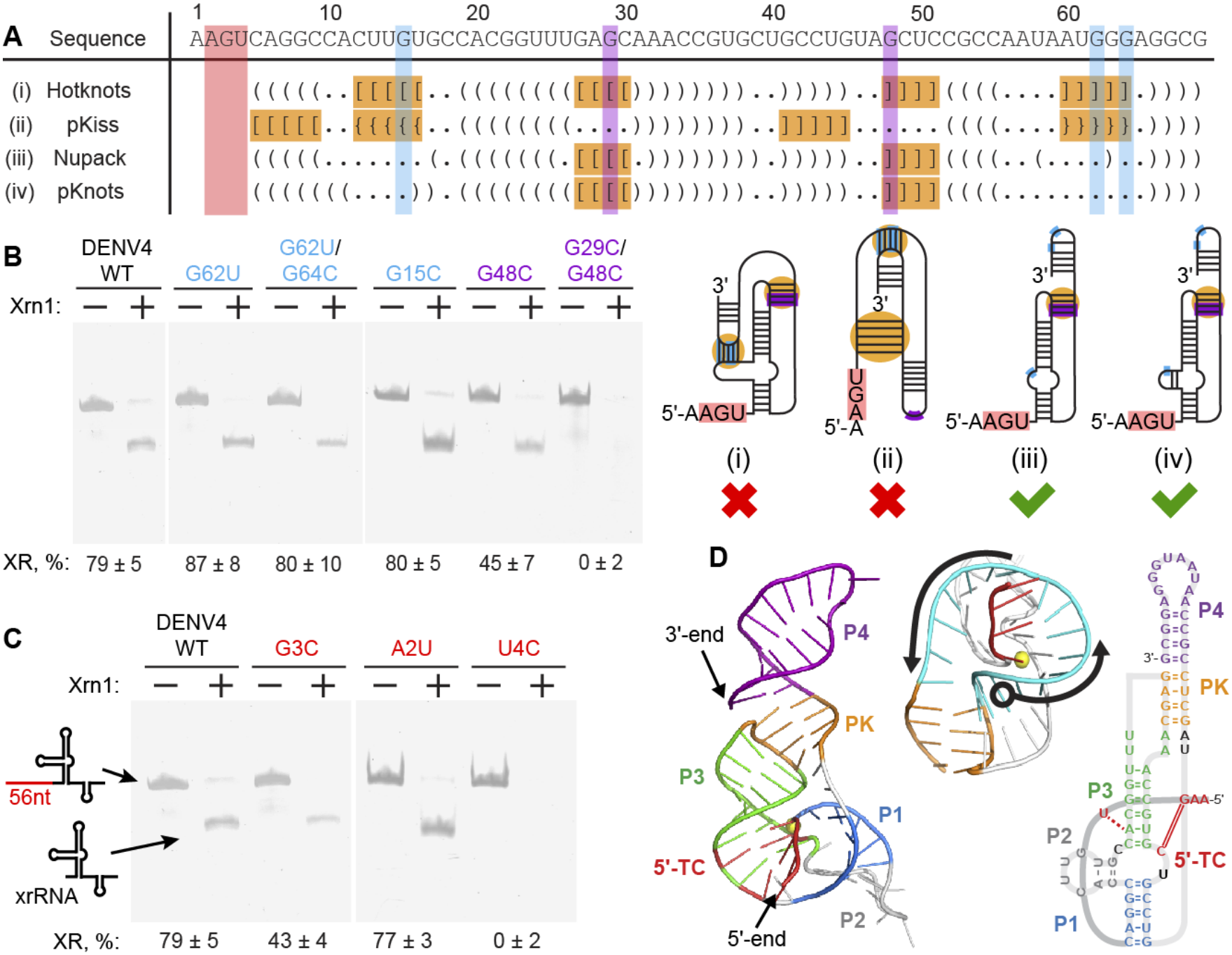
Modeling 3D structure of DENV4 xrRNA. (A) Sequence and potential secondary structures predicted by four structure solvers, shown in dot-bracket notation and cartoons (insets i-iv). Orange: predicted pseudoknots; blue: nucleotides mutated to target pseudoknot interactions; red: nucleotides mutated to target 5′-TC. (B) Representative gels of Xrn1 digestion for pseudoknot mutants. Secondary structures consistent with digestion assay results marked with green checkmarks on panel (A) inset. (C) Representative gels of showing digestion of 5′-TC mutants. (D) Final 3D structural model and 2D representation with colour-coded P1-P4, PK, 5′-TC, and Mg^2+^ ion (yellow). Final structure features knotlike fold with 5′-end (red) threaded through intramolecular ring (cyan) closed by pseudoknot (orange).

MD simulations of these predicted structures found that the base-pairs unique to the prediction arising from the Fig. 4A (iii) model dissociated, whereas all the contacts in the Fig. 4A (iv) model persisted, suggesting that the latter model is the correct one. The 3D structure from the MD simulation showed the same knot-like architecture seen for WNV and other flaviviral xrRNAs, this time with two contacts stabilizing the 5′ end within the ring: A2:U40 and G3:C39 (Fig. S1B). To confirm the presence of these contacts experimentally and test if U4 is also involved in tertiary contacts, as for WNV and ZIKV, we mutated A2, G3, and U4 individually and tested XR levels of these mutants (Fig. 4C).

Whereas breaking G3:C39 with the G3C mutation reduced XR by a factor of ∼2, from 79 ± 5% in WT to 43 ± 4%, indicating that G3 plays an important role in defining XR, the A2U mutation had no effect on XR, implying that it is not present or is irrelevant to XR. Mutating U4 to C to disrupt the potential base triple U4•A20:U37, in contrast, led to complete loss of XR, implying that U4 forms a tertiary contact that is critical to XR. To capture these 5′-TC in the 3D model, we redid the MD simulations starting from structure predictions where the 5′-TC were specified as hard restraints, as above, with and without Mg^2+^. As for the WNV xrRNA, the 5′-TC persisted when Mg^2+^ was present but not in its absence; the final 3D model is shown in Fig. 4D. An overlay of the top 3 cluster centres and the RMSD and RMSF plots are shown in Fig S2B.

### Modeling the DENV2 xrRNA1 structure

Finally, we modelled the structure of the first xrRNA in DENV2, to explore differences between two serotypes of DENV having different numbers of xrRNAs. Each of the 4 predictors suggested different fold architectures: a double-pseudoknot, two distinct single pseudoknots, and a double hairpin without pseudoknots (Fig. 5A). Because yet another single-pseudoknot architecture was previously proposed based on *in vitro* chemical probing experiments [3] and sequence comparisons [56], we included this 2D structure among the possibilities to be tested by mutagenesis and XR measurements.

**Fig. 5:**
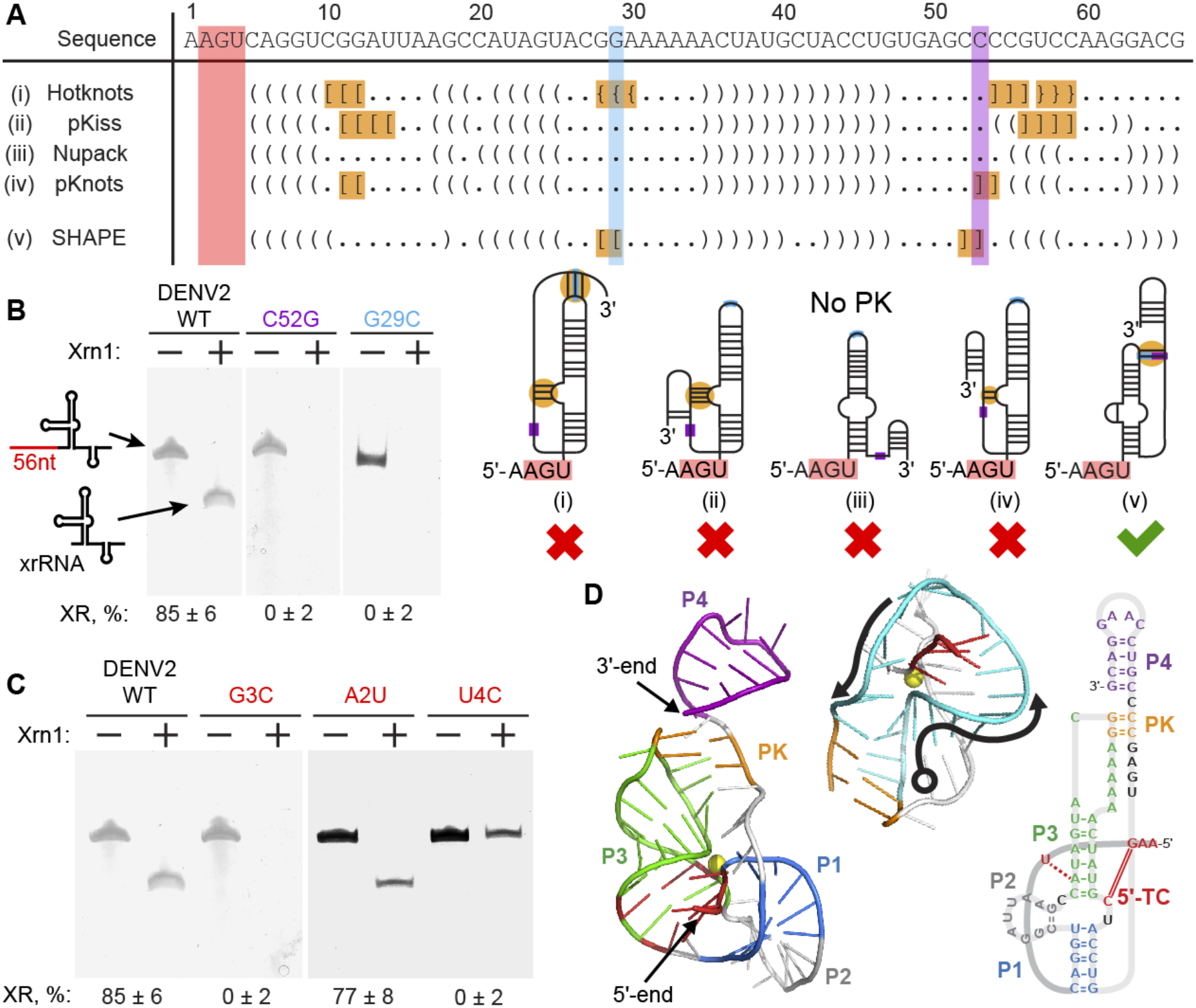
Modeling 3D structure of DENV2 xrRNA1. (A) Sequence and potential secondary structures predicted by four structure solvers plus structure proposed from SHAPE data [3], shown in dot-bracket notation and cartoons (insets i-v). Orange: predicted pseudoknots; blue: nucleotides mutated to target pseudoknot interactions; red: nucleotides mutated to target 5′-TC. (B) Representative gels of Xrn1 digestion for pseudoknot mutants. Secondary structures consistent with digestion assay results marked with green checkmarks on panel (A) inset. (C) Representative gels of showing digestion of 5′-TC mutants. (D) Final 3D structural model and 2D representation with colour-coded P1-P4, PK, 5′-TC, and Mg^2+^ ion (yellow). Final structure features knot-like fold with 5′-end (red) threaded through intramolecular ring (cyan) closed by pseudoknot (orange).

We first mutated G29, which is pseudoknotted in 2 of the 5 models but unpaired in the other, to break the proposed pseudoknot pairings (Fig.5A, blue). G29C abolished XR entirely (Fig. 5B), implying that the models where G29 is unpaired are invalid. To distinguish the two remaining possibilities, we mutated C52 to G (Fig. 5A, purple), which should affect only the model previously proposed in the literature. This mutation again led to complete loss of XR (Fig. 5B), validating the previously proposed pseudoknot, which we used as input for 3D structure prediction by FARFAR2.

The most populated cluster from MD simulations based on the lowest-energy FARFAR2 predictions (Fig. S1C) showed the same threaded knot-like architecture seen in the other examples. Surprisingly, P2 remained stable in the simulations with only 1 bp in the stem, but no contacts stabilizing the 5′ end were seen. Because a lack of 5′-TC seems unlikely based on phylogenetic analysis and SHAPE data [3, 10, 24, 25, 27], as well as the need for such interactions to generate the mechanical rigidity underlying RNase resistance [39, 40], we tested the same 5′-end mutants as for the DENV4 xrRNA. As previously, A2U did not affect XR levels, implying that A2 is unpaired or is not relevant to RNase resistance, whereas U4C abolished XR (Fig. 5C). However, G3C also completely abolished XR, instead of reducing it only by half, implying not just that the G3:C41 pair and U4•A21:U39 triple are present but also that they are each essential for XR on their own, as for ZIKV but unlike for WNV or DENV4. These results suggest that DENV2 xrRNA1 is very sensitive to any changes in the important regions (PK and 5′-TC), since any mutation in those regions completely disrupts the formation of the resistant structure. We refined the 3D structure model to include the 5′ end contacts, again by simulating the xrRNA with and without Mg^2+^. As for WNV and DENV4 xrRNAs, the 5′-TC were unstable without Mg^2+^; the final 3D model is shown in Fig. 5D. An overlay of the top 3 cluster centres and the plots of the RMSD and RMSF values are shown in Fig S2C.

## DISCUSSION

These results show that the WNV and DENV xrRNAs all share the same 3D fold topology, identical to that found previously for other flaviviral xrR-NAs [1, 29-31]: the 5′ end is threaded through a ring formed by a multi-helix junction and closed by a pseudoknot. The same two tertiary contacts held the 5′ end rigidly in place within the ring in all three xrRNAs and were crucial for RNase resistance: a triple involving U4 and a pseudoknot involving G3. The only major difference between the structures was the length of the pseudoknot closing the ring, which varied from 2 bp in DENV2 to 8 bp in WNV.

Considering how these models compare to previous work, we found that the helices forming the junction in our models matched very closely with the results of previous phylogenetic and SHAPE analyses [3, 55, 56]. Turning to the pseudoknot regions of the xrRNAs, the results we found for both DENV variants are identical to the pseudoknots proposed based on chemical probing and sequence comparisons, with four and two base-pairs in the pseudoknots of DENV4 and DENV2, respectively [3, 55, 56, 60]. However, the pseudoknot we found for the WNV xrRNA differed from that proposed based on computational and low-resolution experimental predictions, which suggested it consisted of 6 bp [38, 55, 56] as opposed to the 8 pairs in our model. Such a 6-bp PK is inconsistent with the results of the XR assays: breaking 4 or 5 of the pairs in a 6-bp PK would be expected to abolish XR based on past results from the ZIKV xrRNA1 (where breaking 2 of 5 pairs did so [40]), yet breaking 4 pairs with the mutations G32C/U33A/G35C/G37C did not noticeably reduce XR (Fig. S3), and breaking 5 pairs in the G35C/A36U/G56C/A60U/A61U/C62G mutant only reduced XR activity by a third.

The fact that the same two 5′ end tertiary contacts were present in all the xrRNAs, identical to the contacts present in the ZIKV xrRNA1 [29], is perhaps not surprising, given that these contacts are essential for anchoring the 5′ end to the ring and rigidifying the structure sufficiently to generate RNase resistance [39, 40]. Curiously, however, the A2:U pair implied by conservation and present in some 3D models (*e*.*g*. DENV4) does not seem to contribute to RNase resistance, in contrast to the G3:C and U4•A:U contacts. The observation from the DENV4 xrRNA that XR activity depends more strongly on the U4•A:U base triple than the G3:C pair is consistent with previous results from the ZIKV xrRNA1 that U4•A:U rigidifies the structure more than does G3:C [40]. As with the ZIKV xrRNA1, Mg^2+^ ions appear to be essential to maintaining these contacts, and thus to the mechanical rigidity central to XR activity [39].

The mutants studied here also extend our understanding of the relative importance of the 5′-TC and PK contacts for supporting XR activity. Generally, no single PK contact was essential for high XR (with the exception of DENV2), whereas each 5′-TC was individually important (with the exception of WNV). This hierarchy of importance makes sense given that the RNase pulls directly on the 5′ end: the 5′-TC become the primary load-bearing contacts in the structure, distributing mechanical stress to the ring as the xrRNA resists digestion, whereas the PK interactions play a supporting role in maintaining the structure of the ring [40]. The PK interactions only become as important as the stabilizing 5′-TC when the pseudoknot is very short, as in DENV2, where the loss of any one contact is enough to destabilize the pseudoknot and open the ring. Conversely, in the WNV xrRNA, the very long PK region may collectively contribute sufficient rigidity to the ring on its own such that deleting any one of the 5′-TC does not reduce RNase resistance.

This work illustrates the effectiveness of the approach we adopt here, integrating secondary structure prediction, site-directed mutagenesis, enzymatic assays, and MD modeling to solve for structures of xrRNAs that are challenging to obtain from standard high-resolution methods [61]. Computational methods alone frequently fail to predict crucial tertiary interactions within highly structured functional RNAs, even using advanced machine-learning platforms [62], highlighting the need for additional experimental inputs. This problem is resolved here by incorporating iterative refinement of modeling via assays of functional activity within the workflow.

However, our approach does have important limitations. First, the secondary-structure predictors are constrained by simplified energy models, limited thermodynamic parameter sets, and assumptions of equilibrium folding, which likely explain why none of the platforms predicted the correct 2-bp pseudoknot for DENV2. Second, any inaccuracies in secondary-structure prediction cascade directly into the 3D modeling. Third, the workflow cannot discriminate easily between predicted secondary structures that differ in regions that do not affect activity (*e*.g., P2 or P4 stem-loops here), as seen for DENV4. Lastly, Mg^2+^ ion positions can only be modelled by similarity to known structures. Mg^2+^ ions are known to be important for stabilizing 5′-TC in ZIKV xrRNA1 [39] and closing the pseudoknot in WNV xrRNA1 [63], but MD simulations are an unreliable guide to the number and position of these ions, and high-resolution experimental methods are needed to confirm their location more reliably.

We note that the presence in each of the structures solved here of the knot-like fold observed previously [1, 29-31] suggests that this fold topology is likely to be universal among all *Flaviviridae* xrRNAs, which are also therefore likely to share the same mechanically based resistance to 5′-to-3′ processive degradation enzymes. The fact that xrRNAs from other virus families have also revealed a knot-like fold with very similar features [7] suggests that this mechanism of RNase resistance may well be universal across evolutionarily distant virus families of viruses, a hypothesis that will need to be tested in future studies of the mechanical properties of other xrRNAs. Finally, although our analysis here focused on flaviviral xrRNAs, the same strategy of combining computational modeling with functional assays could be applied to other structured RNAs which function depends on complex tertiary interactions, including regulatory RNAs and structured viral elements in unrelated families.

## Acknowledgements

This work was supported by the Canadian Institutes of Health Research (grant reference number PJT-183591). SN acknowledges fellowship support from the Canadian Institutes of Health Research. Computational resources were provided by the Digital Research Alliance of Canada and Prairies DRI Group.

## Author contributions

Conceptualization: MTW, DZ, RVS; Resources: DZ, SM, IM; Investigation: DZ, SM, RVS; Formal analysis: DZ, RVS; Supervision: MTW; Funding acquisition: MTW; Visualization: DZ, RVS, MTW; Writing—original draft: DZ, RVS; Writing—review & editing: DZ, SM, RVS, IM, MTW.

## Supporting Information

**Table S1:**
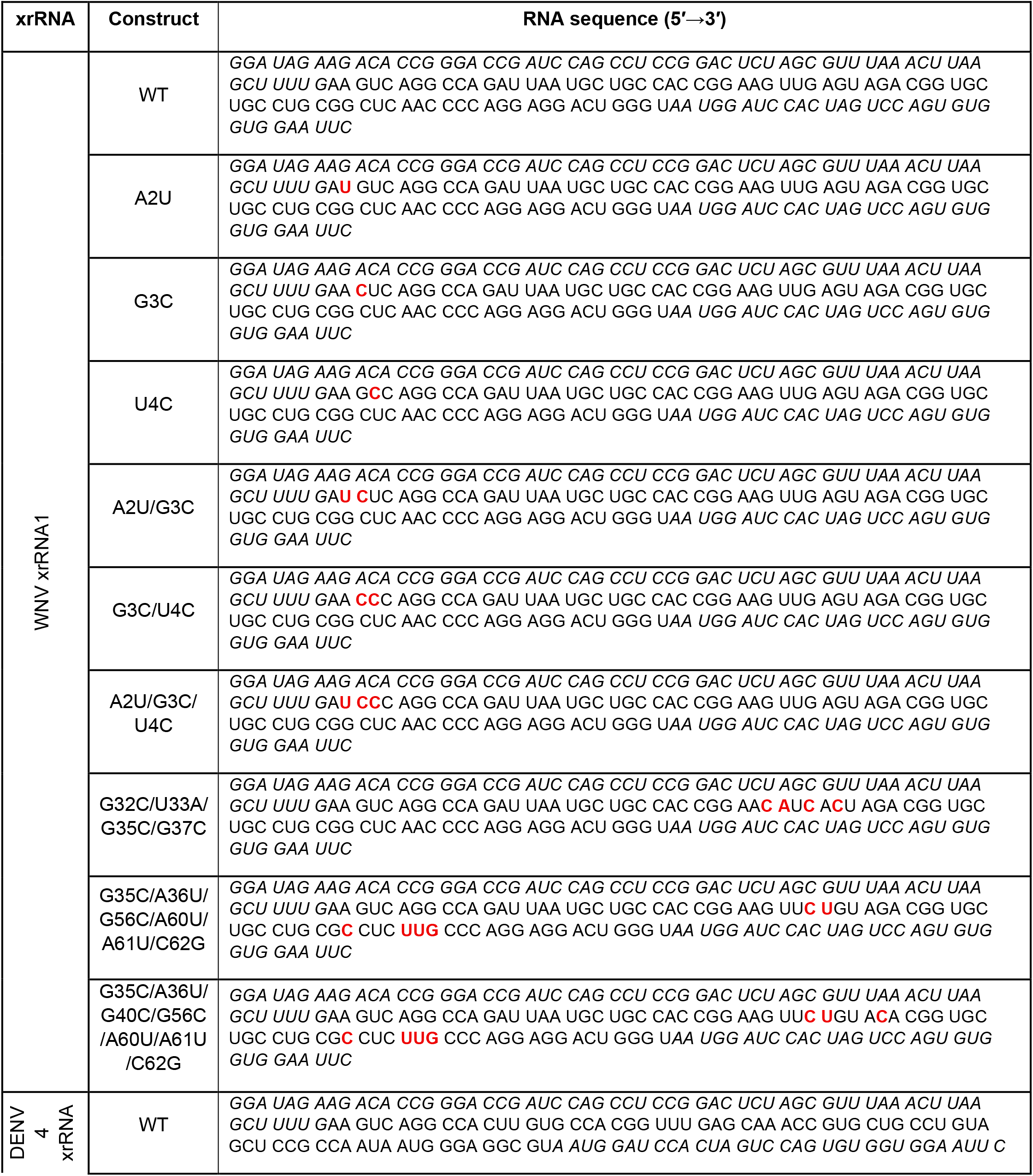
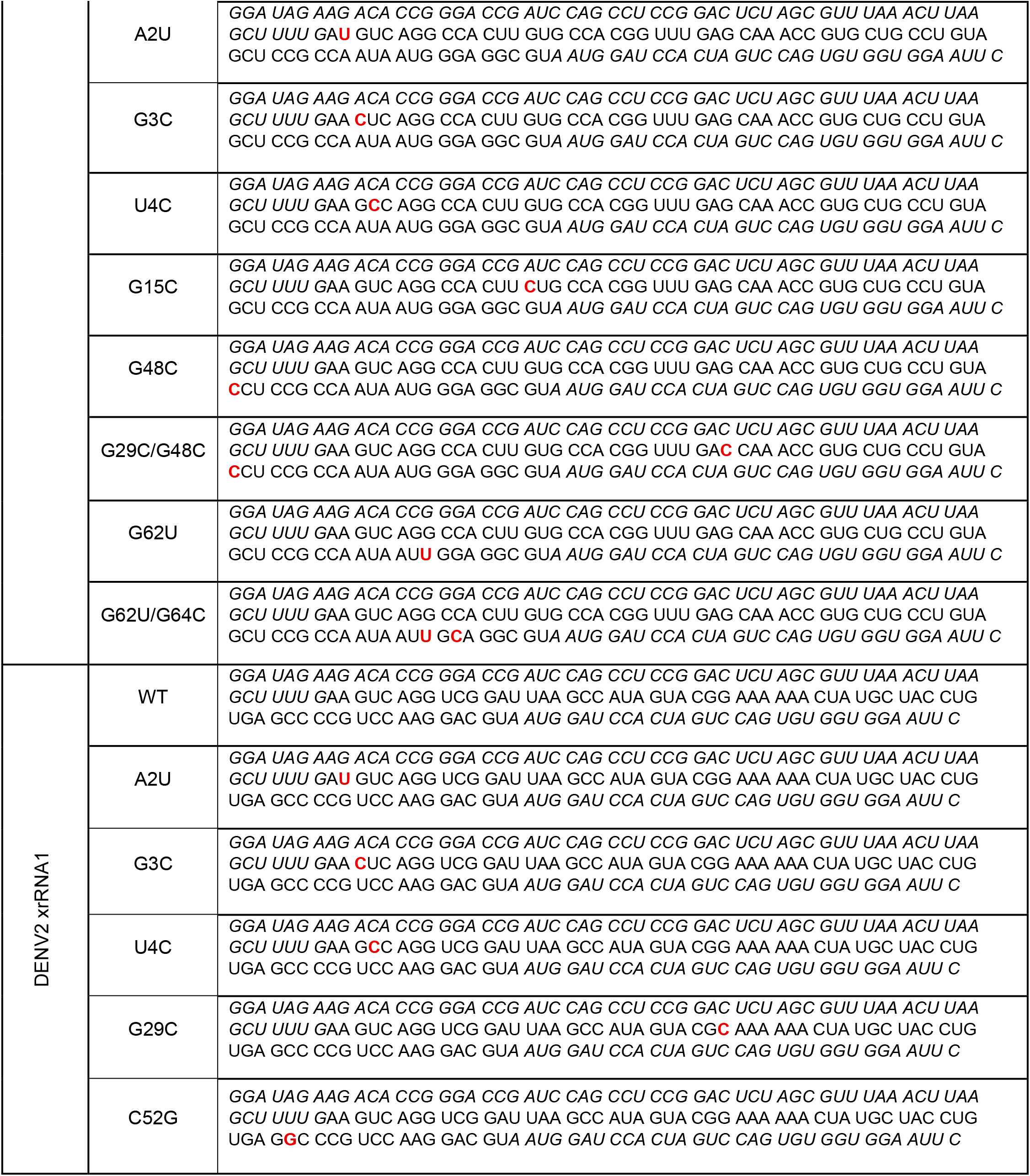
xrRNA sequences. Sequences of flaviviral xrRNA constructs used for Xrn1 digestion assay. Mutations shown in red, leader and downstream sequences in italics.

**Fig S1:**
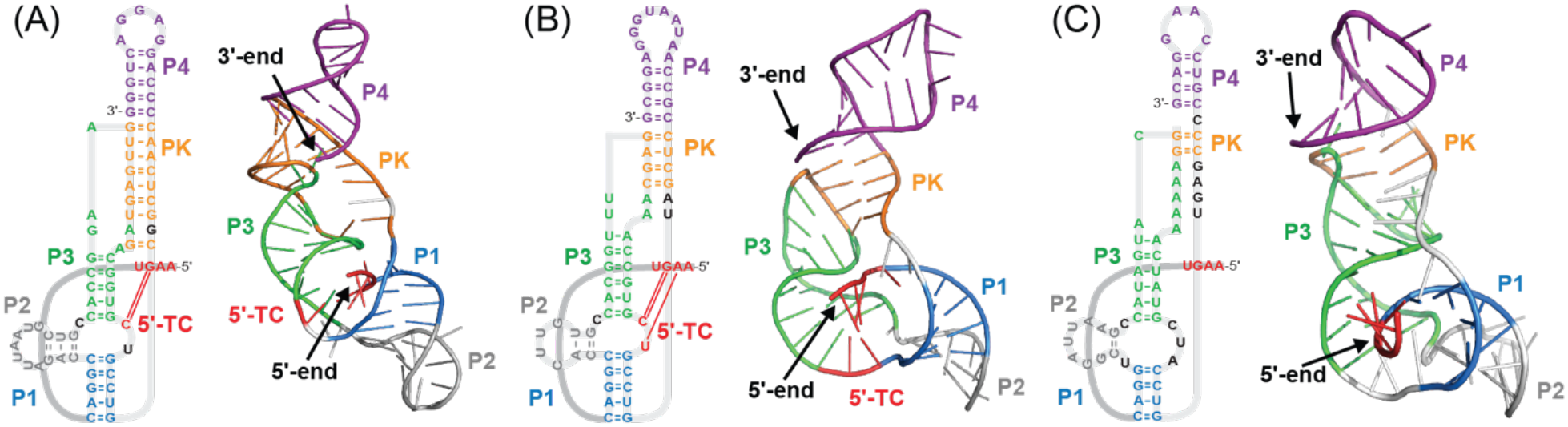
Initial 3D models of flaviviral xrRNAs. After completing the first round of MD simulations starting with the lowest-energy prediction from FARFAR2 for each of the 2D models, the most populated clusters converged on the represented 3D structures of WNV xrRNA1 (A), DENV4 xrRNA (B), and DENV2 xrRNA1 (C) with colour-coded P1-P4, PK, and 5′-TC. All of the predictions show a formation of a ring-like topology with different number of stabilizing 5′-TC: from 2 in DENV4 to none in DENV2.

**Fig S2:**
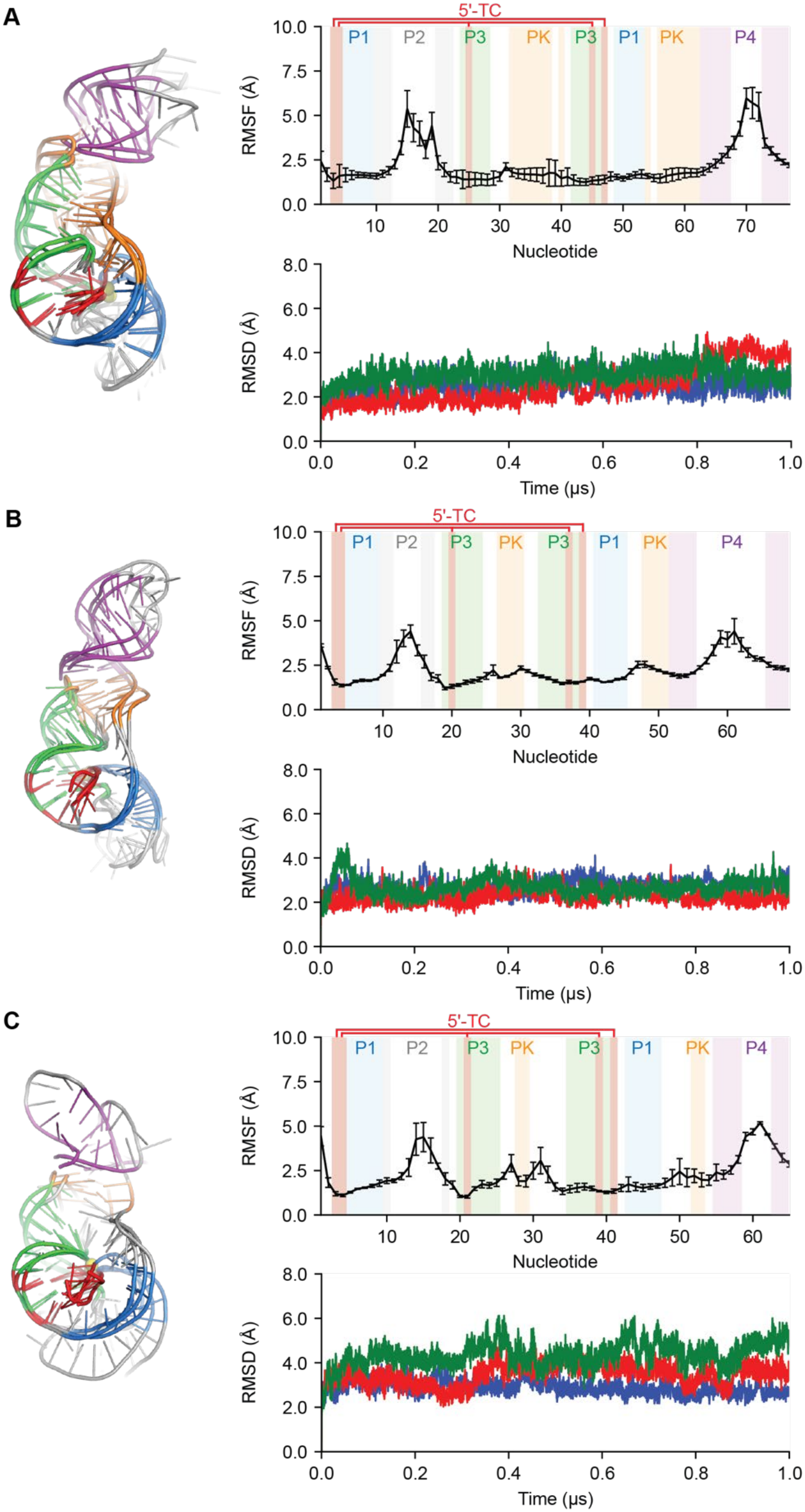
Final 3D models from MD simulations. Overlay of the final 3D structures of the three most populated clusters from MD simulations for WNV xrRNA1 (A), DENV4 xrRNA (B), and DENV2 xrRNA1. Top inset: mean RMSF values for each nucleotide with error bars presenting S.E.M. with colour-coded P1-P4, PK, and 5′-TC. Bottom inset: RMSD vs. time plots for three independent simulations.

**Fig S3:**
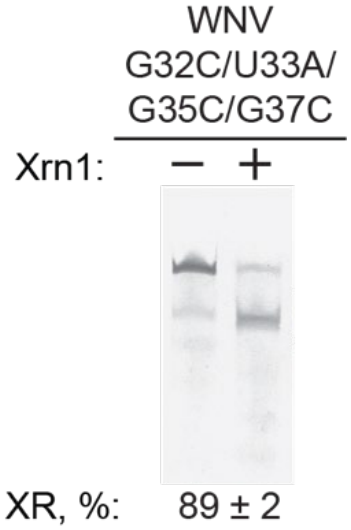
Xrn1 Assay of WNV xrRNA1 G32C/U33A/G35C/G37C mutant. Breaking 4 base pairs in the 8bp-long pseudoknot of WNV xrRNA1 does not alter its XR level compared to wild-type.

